# The Recovery–Burden Index for Assessing *β*-Cell Function from OGTT Glucose Profiles

**DOI:** 10.64898/2026.05.17.725721

**Authors:** Ren Zhang

## Abstract

Disposition index (DI) is an informative measure of *β*-cell function adjusted for insulin resistance, but its assessment is procedurally demanding, requiring dynamic testing with timed sampling and insulin or C-peptide–based estimation of insulin sensitivity and secretion. A simple glucose-only metric derived from the oral glucose tolerance test (OGTT) could provide a practical approach to estimating DI. We developed the Recovery–Burden Index (RBI), a glucose-only geometric metric that quantifies post-peak glucose recovery relative to total glucose excursion during OGTT. Using densely sampled venous OGTT profiles with measured DI, RBI was evaluated for prediction of continuous DI by leave-one-out (LOO) cross-validated *R*^*2*^ and for discrimination of DI-defined *β*-cell dysfunction by AUROC. Performance was compared with conventional glycemic metrics. RBI predicted continuous DI more accurately than conventional glycemic metrics, with LOO *R*^*2*^ of 0.43, Pearson *r =* 0.70, and Spearman *ρ =* 0.75. RBI30–180 performed similarly, with cross-validated *R*^*2*^ of 0.42, Pearson *r =* 0.72, and Spearman *ρ =* 0.75. RBI also discriminated DI-defined *β*-cell dysfunction, with AUROC values of 0.90 for RBI and 0.91 for RBI30–180. Reduced sampling schedules preserved much of the RBI signal, whereas truncation at 120 min attenuated continuous DI prediction, supporting the contribution of late recovery-phase information. RBI extracts β-cell–relevant information from the OGTT glucose profile using a single transparent glucose-only index. These findings highlight post-peak recovery as a key feature for estimating DI-associated β-cell compensation and support further validation of RBI in extended or CGM-augmented OGTT settings.

## Introduction

Prediabetes and type 2 diabetes are diagnosed primarily by elevated glucose levels, including fasting glucose, 2-hour OGTT glucose, and HbA1c [1-3]. These measurements define dysglycemia but do not directly quantify *β*-cell function, a key determinant of glucose tolerance and diabetes progression. *β*-cell function is difficult to interpret from insulin secretion alone because secretion must be matched to the prevailing degree of insulin resistance [4, 5]. When insulin resistance is greater, a higher insulin secretory response is required to maintain normal glucose levels. Thus, the same insulin secretory response may represent adequate compensation in an insulin-sensitive individual but insufficient compensation in an insulin-resistant individual.

Disposition index (DI) addresses this limitation by evaluating insulin secretion relative to insulin sensitivity [5-7]. Because insulin secretion normally increases as insulin sensitivity declines, DI reflects whether *β*-cell insulin secretory responses are appropriate for the prevailing degree of insulin resistance. DI is therefore a physiologically informative measure of *β*-cell compensation and is widely used in metabolic research [8, 9]. However, quantitative β-cell function assessment remains procedurally demanding. Clamp- and IVGTT-based methods provide detailed physiological information, and OGTT-or meal-test–based modeling improves feasibility for larger studies, but these approaches still require timed insulin or C-peptide sampling and specialized analysis [10]. DI assessment further requires protocol-dependent estimation of insulin sensitivity and insulin secretion, which may involve OGTT-based indices or modeling, C-peptide deconvolution, or independent measures of insulin sensitivity [11, 12]. This complexity limits the feasibility of DI assessment in routine clinical care and large-scale epidemiological studies. A simple glucose-only metric that captures DI-related information from the OGTT curve would therefore be valuable for metabolic phenotyping.

The OGTT provides a dynamic glucose challenge and therefore contains more information than fasting or 2-hour glucose alone. However, routine interpretation of the OGTT typically relies on selected timepoint concentrations, fasting, 1-hour, or 2-hour glucose, or on summary exposure measures such as total glucose AUC. These metrics are clinically useful but incompletely describe the structure of the glucose response, including peak timing, post-peak recovery, and residual glycemic burden. Dynamic glucose profiles contain physiologically relevant information beyond fasting or 2-hour glucose. OGTT curve patterns have been associated with insulin sensitivity, β-cell function, and future diabetes risk [13], and recent model-based or machine-learning approaches have used glucose dynamics to estimate DI or predict metabolic subphenotypes [12, 13]. These findings suggest that β-cell–relevant information is embedded in the glucose response curve, but they also raise a practical question: can this information be captured by a simpler, transparent, single-number metric?

We developed the Recovery–Burden Index (RBI) as a glucose-only geometric metric that summarizes post-peak recovery relative to total glucose excursion. RBI was designed to quantify the balance between resolution of the glucose excursion after the peak and residual post-peak burden, a feature not directly measured by fasting glucose, 2-hour glucose, HbA1c, or total glucose AUC. In this study, we evaluated whether RBI predicts continuous DI and discriminates DI-defined *β*-cell dysfunction in a cohort with densely sampled venous OGTT profiles and measured DI. We compared RBI with conventional glycemic metrics and used representative OGTT profiles to illustrate the recovery patterns reflected by the index.

## Materials and Methods

### Data source and OGTT measurements

This study was a secondary analysis of the publicly available Stanford metabolic subphenotyping dataset [13]. We analyzed the initial deeply phenotyped cohort of 32 participants who completed standardized metabolic testing in the Stanford Clinical and Translational Research Unit and had both reference DI measurements and high-resolution OGTT glucose profiles available. Although the source study included additional participants, the full paired dataset required for the present DI-based analysis was not available for all participants in the public data release. Therefore, the present analysis was restricted to the n = 32 participants with available DI and 16-timepoint OGTT glucose data. Participants underwent a 75-g, 3-h oral glucose tolerance test after an overnight fast. Plasma glucose was measured at −10, 0, 10, 15, 20, 30, 40, 50, 60, 75, 90, 105, 120, 135, 150, and 180 min. For the present analysis, only the 0–180 min venous glucose curve was used. DI values were obtained from the source dataset. In the source study, DI was calculated from C-peptide deconvolution-derived insulin secretion adjusted for insulin resistance assessed by steady-state plasma glucose. β-cell dysfunction was defined as DI < 1.58, following the source-study cutoff.

### Recovery–Burden Index

Let *G*(t) denote plasma glucose concentration at time *t* during the analyzed OGTT interval, with *G*_min_ and *G*_max_ denoting the minimum and maximum glucose values, respectively, and *t*_max_ denoting the time of peak glucose. The glucose-excursion area was calculated as the area between the glucose curve and *G*_min_:

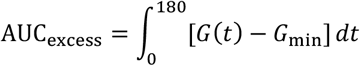

The post-peak area above the glucose curve and below *G*_max_, *AAC*_*a*_, was calculated from *t*_max_ to 180 min:

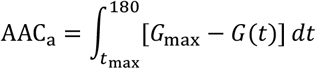

The Recovery–Burden Index (RBI) was defined as:

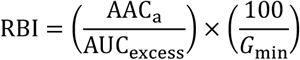

RBI is therefore a recovery-to-burden measure: the numerator quantifies post-peak recovery, whereas the denominator quantifies total glucose-excursion burden above the participant-specific minimum. Higher RBI indicates greater post-peak recovery relative to total glucose-excursion burden, whereas lower RBI indicates persistent post-peak hyperglycemia. The 100/G_min_ term provides a subject-specific glucose-level normalization, with 100 serving as a unit-scaling constant for glucose expressed in mg/dL. All RBI values in this study were calculated using glucose in mg/dL. All areas were calculated using trapezoidal integration. RBI was calculated over the 0–180 min OGTT interval. A secondary version, RBI30–180, was calculated after restricting the glucose curve to 30–180 min to focus on the postchallenge recovery phase.

### Comparator glycemic and clinical metrics

RBI was compared with conventional OGTT-derived and clinical glycemic metrics. OGTT time-point metrics included fasting plasma glucose, *G*_60_, and *G*_120_, defined as plasma glucose concentrations at 0, 60, and 120 min, respectively. Total OGTT glucose exposure was quantified by trapezoidal integration of the glucose curve from 0 to 180 min:

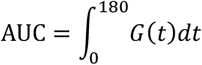

HbA1c was included as a standard clinical marker of chronic glycemia. Comparator metrics were evaluated individually using the same DI prediction and DI-defined *β*-cell dysfunction analyses as RBI.

### Evaluation of RBI for predicting disposition index and DI-defined *β*-cell dysfunction

The primary analysis evaluated whether RBI and comparator metrics predicted continuous DI, a quantitative measure of *β*-cell function adjusted for insulin resistance. Each metric was evaluated individually as a single predictor in a linear regression model with DI as the outcome. Because of the small sample size, predictive performance was assessed using leave-one-out cross-validation, in which each participant was held out once and predicted from a model fit to the remaining participants. Cross-validated *R*^*2*^ was used as the primary criterion for ranking metrics, because the main objective was prediction of continuous *β*-cell function rather than classification alone. For each metric, we also calculated ordinary least-squares *R*^*2*^ from the full cohort as an apparent in-sample measure of model fit. Pearson correlation was used to quantify linear association with DI, and Spearman correlation was used to quantify rank-based association.

As a secondary analysis, we evaluated binary discrimination of DI-defined *β*-cell dysfunction. *β*-cell dysfunction was defined using the source-study cutoff of *DI* < 1.58. Receiver operating characteristic curves were generated for each metric, and area under the receiver operating characteristic curve (AUROC) was calculated. For metrics positively associated with preserved *β*-cell function, including RBI, the score direction was reversed where necessary so that lower values indicated greater probability of *β*-cell dysfunction. For conventional glucose-burden metrics, higher values generally indicated greater probability of *β*-cell dysfunction. AUROC values were therefore oriented consistently, such that higher AUROC always indicated better discrimination of DI-defined *β*-cell dysfunction.

### Sensitivity analyses for sampling density and analysis-window duration

To evaluate whether RBI performance depended on dense OGTT sampling, RBI was recalculated after downsampling the 0–180 min glucose curve to reduced sampling schedules. For each reduced schedule, G_min_, G_max_, AUC_excess_, and AAC_a_ were recalculated using only the retained timepoints, and areas were estimated by trapezoidal integration over the reduced sampling grid. To evaluate the contribution of OGTT analysis-window duration, RBI was recalculated using windows beginning at 0 or 30 min and ending at 120, 150, or 180 min. For each window, all RBI components were recalculated using only glucose values within that interval. Each downsampled or window-restricted RBI version was evaluated using the same continuous DI prediction and DI-defined β-cell dysfunction analyses as the primary RBI.

### Statistical analysis and software

All areas under or above glucose curves were calculated using trapezoidal integration over the observed OGTT sampling times. For continuous DI prediction, each metric was evaluated in a separate linear regression model. Leave-one-out cross-validation was used to estimate out-of-sample predictive performance, reported as cross-validated *R*^*2*^. Ordinary least-squares *R*^*2*^ was also reported as the apparent in-sample model fit. Associations between each metric and DI were assessed using Pearson correlation for linear association and Spearman correlation for rank-based association. Differences in RBI between participants with DI < 1.58 and DI ≥ 1.58 were assessed using the two-sided Mann–Whitney U test. For classification of DI-defined *β*-cell dysfunction, receiver operating characteristic curves were generated using the metric values as prediction scores, and AUROC values were calculated. For metrics inversely associated with DI, prediction direction was oriented so that higher AUROC consistently indicated better discrimination of *β*-cell dysfunction. All statistical tests were two-sided, and *p*< 0.05 was considered statistically significant. All analyses were performed in Python using pandas, NumPy, SciPy, scikit-learn, statsmodels, and matplotlib.

## Results

### Dissection of OGTT glucose profiles and construction of the Recovery–Burden Index

We first decomposed each venous OGTT glucose profile into geometric components reflecting glucose excursion and post-peak recovery (Fig. 1A). For each participant, the minimum and maximum glucose concentrations during the analyzed OGTT interval were identified. Total glucose excursion was quantified as the excess area under the curve, defined as the area between the glucose curve and the participant-specific minimum glucose concentration. This area represents the overall glucose burden above the participant-specific minimum. The excess area under the curve can be divided into pre-peak and post-peak components, corresponding to the rising glucose burden before peak glucose and the residual glucose burden after peak glucose. The post-peak area above the curve was calculated as the area between the maximum glucose concentration and the observed glucose curve from the time of peak glucose to 180 min; this area is large when glucose falls rapidly after the peak and small when glucose remains persistently elevated (Fig. 1A).

**Figure 1.**
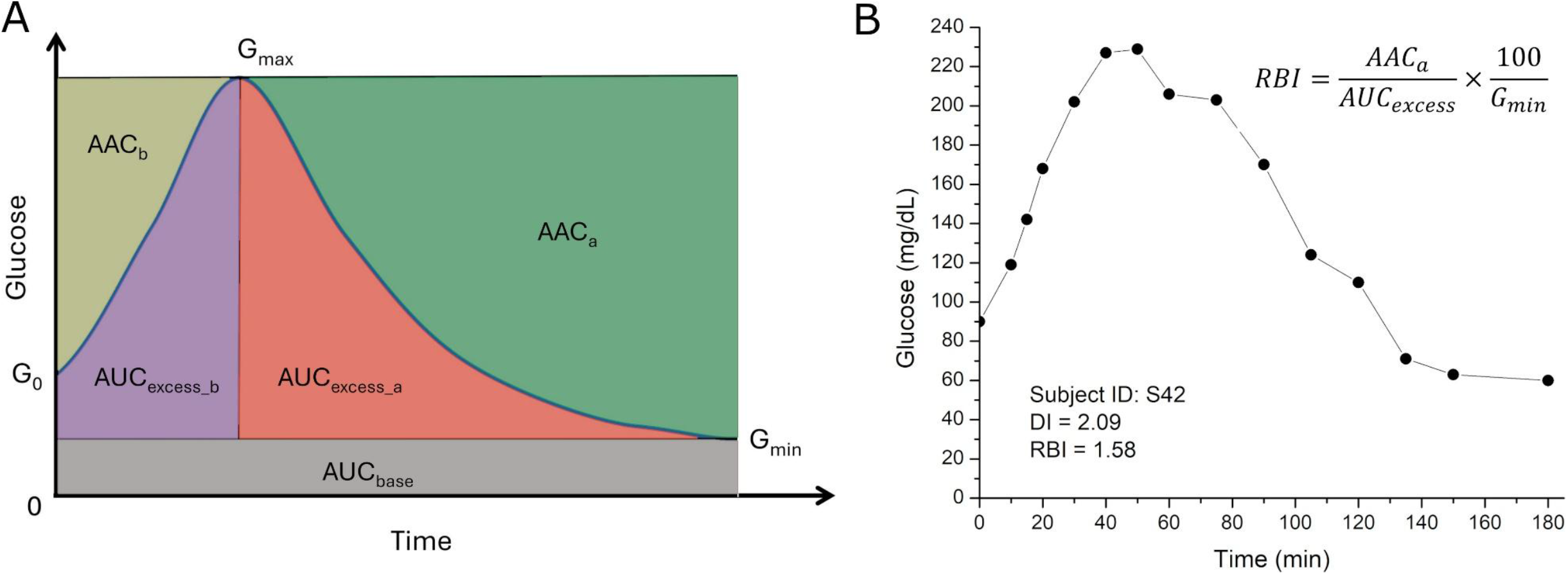
Definition and representative example of the Recovery–Burden Index (RBI). **(A)** Schematic OGTT glucose profile showing G_min_, G_max_, AAC_a_, and the pre- and post-peak components of glucose-excursion area. AUC_excess_ represents the sum of AUC_excess_b_ and AUC_excess_a_. **(B)** Representative venous OGTT profile from subject S42 with preserved *β*-cell function. RBI is calculated as post-peak AAC_a_ divided by AUC_excess_, adjusted by 100 divided by G_min_. DI, disposition index; RBI, Recovery–Burden Index.

RBI was constructed as a recovery-to-burden measure of the OGTT glucose profile. The numerator, AAC_a_, quantifies the post-peak area above the glucose curve; it is large when glucose declines rapidly after the peak and small when post-peak hyperglycemia persists. The denominator, AUC_excess_, quantifies the total glucose excursion relative to the participant-specific minimum glucose level. Thus, AAC_a_/AUC_excess_ measures post-peak recovery relative to the overall glucose-excursion burden (Fig. 1B).

RBI further includes a 100/G_min_ normalization to incorporate the glycemic background on which the excursion occurs. For the same recovery-to-burden geometry, a higher minimum glucose level indicates a higher glucose profile throughout the analyzed interval and therefore less favorable glycemic regulation. The constant 100 serves as a scaling factor that anchors the index to a typical normoglycemic glucose value and keeps RBI values on a convenient numerical scale. RBI was calculated over the full 0–180 min OGTT interval. RBI30–180 was evaluated as a secondary version focused on the postchallenge recovery interval, where recovery dynamics may be more strongly represented and less influenced by the initial glucose rise. Higher RBI indicates greater recovery relative to burden, whereas lower RBI indicates persistent post-peak hyperglycemia.

### RBI predicts continuous disposition index

We next evaluated whether RBI predicts disposition index in the 32 participants with measured DI. DI is a continuous measure of *β*-cell function adjusted for insulin resistance. Because the cohort size was small, leave-one-out cross-validated *R*^*2*^ was used as the primary benchmark of predictive performance. This metric estimates how well each single-predictor model generalizes to held-out participants. Ordinary least-squares *R*^*2*^ was also reported as the apparent in-sample model fit. Pearson correlation was used to assess linear association with DI, whereas Spearman correlation was used to assess rank-based association less dependent on linearity. RBI showed the strongest out-of-sample prediction among the evaluated metrics, with a leave-one-out cross-validated *R*^*2*^ of 0.43 and an ordinary least-squares *R*^*2*^ of 0.49. RBI was positively associated with DI by both Pearson correlation (*r =* 0.70, *p =* 7.2×10^−6^) and Spearman correlation (*ρ =* 0.75, *p =* 6.7×10^−7^), indicating that higher RBI was associated with better preserved *β*-cell function. The 30–180 min version of RBI performed similarly, with a cross-validated *R*^*2*^ of 0.42 and an ordinary least-squares *R*^*2*^ of 0.52. RBI_30–180_ was also strongly associated with DI by Pearson correlation (*r =* 0.72, *p =* 3.7×10^−6^) and Spearman correlation (*ρ =* 0.75, *p =* 8.5×10^−7^).

Conventional glycemic metrics showed weaker prediction of continuous DI. G120 was the strongest conventional comparator, but its cross-validated *R*^*2*^ was lower than RBI (0.25 versus 0.43). Fasting plasma glucose and total glucose AUC each had cross-validated *R*^*2*^ values of 0.15, followed by G60 at 0.11 and HbA1c at 0.03. These conventional metrics were inversely associated with DI, consistent with higher glucose burden reflecting poorer *β*-cell function, but their predictive performance was lower than that of RBI. These results indicate that RBI captures DI-relevant recovery information in the OGTT glucose profile beyond standard glucose concentrations, total glucose exposure, and chronic glycemia (Table 1).

**Table 1.** Performance of RBI and comparator metrics for predicting disposition index.

| **Metric** | **LOO $R^2$** | **OLS $R^2$** | **Pearson $r$** | **Pearson $p$** | **Spearman $\rho$** | **Spearman $p$** | **AUROC** |
| --- | --- | --- | --- | --- | --- | --- | --- |
| RBI | 0.43 | 0.49 | 0.70 | 7.2E-06 | 0.75 | 6.7E-07 | 0.90 |
| RBI<sub>30-180</sub> | 0.42 | 0.52 | 0.72 | 3.7E-06 | 0.75 | 8.5E-07 | 0.91 |
| G120 | 0.25 | 0.35 | -0.59 | 4.1E-04 | -0.55 | 1.2E-03 | 0.79 |
| FPG | 0.15 | 0.23 | -0.48 | 5.8E-03 | -0.46 | 7.8E-03 | 0.67 |
| AUC | 0.15 | 0.26 | -0.51 | 3.0E-03 | -0.44 | 1.2E-02 | 0.70 |
| G60 | 0.11 | 0.23 | -0.47 | 6.1E-03 | -0.40 | 2.4E-02 | 0.70 |
| HbA1c | 0.03 | 0.12 | -0.35 | 4.9E-02 | -0.41 | 1.9E-02 | 0.69 |

RBI was calculated as $AAC_a$ divided by $AUC_{\text{excess}}$ , adjusted by 100 divided by $G_{\text{min}}$ . RBI<sub>30-180</sub> indicates the OGTT interval used for calculation. Metrics were ranked by leave-one-out cross-validated $R^2$ , the primary criterion for continuous DI prediction. AUROC was calculated for DI-defined β-cell dysfunction, defined as $DI < 1.58$ . For conventional glucose-burden metrics that are inversely associated with DI, the AUROC direction was oriented so that higher AUROC indicates better discrimination. Analysis included 32 participants with measured DI.

### RBI discriminates DI-defined *β*-cell dysfunction

We next evaluated whether RBI discriminates participants with DI-defined *β*-cell dysfunction. Using the source-study cutoff, *β*-cell dysfunction was defined as DI <1.58. Because this analysis converted continuous DI into a binary outcome, it was treated as a secondary analysis. Receiver operating characteristic curves were generated for each metric, and AUROC was used to quantify discrimination. AUROC values were oriented so that higher values consistently indicated better discrimination of DI-defined *β*-cell dysfunction.

RBI values were significantly lower in participants with DI < 1.58 than in those with DI ≥ 1.58, consistent with impaired post-peak glucose recovery in participants with reduced *β*-cell function (Mann–Whitney U test, *p*< 0.001; Fig. 2A). Formal discrimination analysis showed that RBI achieved an AUROC of 0.90 for identifying DI-defined *β*-cell dysfunction. The 30–180 min version performed similarly, with an AUROC of 0.91. In comparison, conventional glycemic metrics showed lower discrimination, including G120 with an AUROC of 0.79, total glucose AUC and G60 with AUROC values of 0.70, HbA1c with an AUROC of 0.69, and fasting plasma glucose with an AUROC of 0.67. These findings indicate that reduced RBI identifies participants with DI-defined *β*-cell dysfunction more effectively than standard glucose concentrations, total OGTT glucose exposure, or HbA1c (Fig. 2B, Table 1).

**Figure 2.**
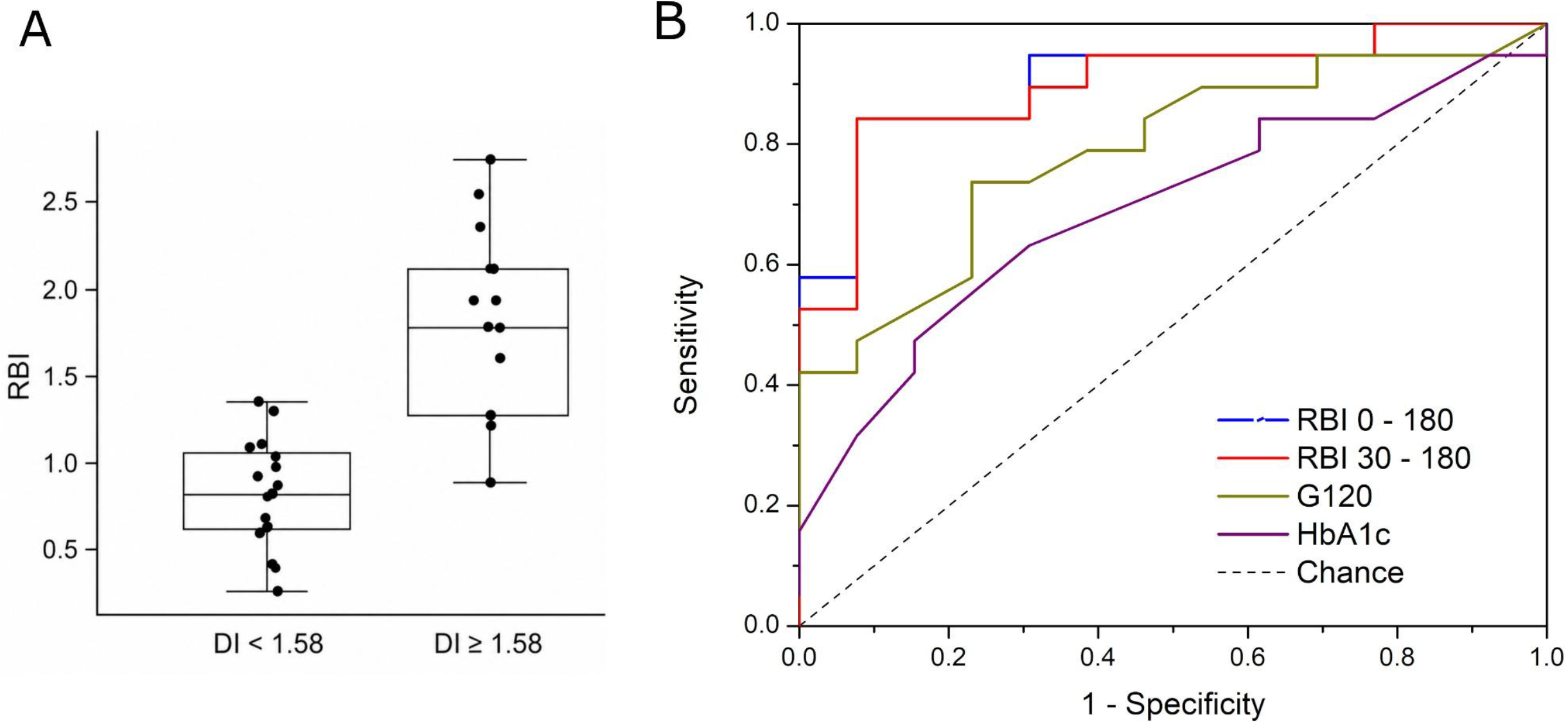
RBI discriminates DI-defined *β*-cell dysfunction. **(A)** RBI values in participants with DI < 1.58 versus DI ≥ 1.58, calculated from the 0–180 min OGTT interval. Each dot represents one participant. The horizontal line inside each box indicates the median, the box indicates the interquartile range, and whiskers indicate the range of values. RBI values were significantly lower in participants with DI < 1.58 than in those with DI ≥ 1.58 (Mann–Whitney U test, *p <* 0.001). **(B)** Standard ROC curves for classifying DI-defined *β*-cell dysfunction, defined as DI < 1.58. RBI was evaluated using both 0–180 min and 30–180 min OGTT intervals and compared with G120 and HbA1c. The dashed diagonal line indicates chance-level discrimination.

### Representative OGTT profiles illustrate recovery-dominant and burden-dominant patterns

To illustrate the biological interpretation of RBI, we compared two representative OGTT profiles with broadly similar fasting and peak glucose levels but markedly different post-peak recovery (Fig. 3). This comparison was selected to highlight that similar early or peak glycemic responses can diverge substantially during the recovery phase. Subject S06 showed a recovery-dominant profile, characterized by rapid glucose decline after the peak, a large post-peak area above the curve, high RBI, and preserved disposition index. In contrast, subject S26 showed a burden-dominant profile, characterized by persistent post-peak hyperglycemia, a smaller post-peak area above the curve, low RBI, and DI-defined *β*-cell dysfunction. These paired profiles show that RBI captures recovery dynamics that are not apparent from fasting or peak glucose alone. Under broadly comparable fasting and peak glucose conditions, greater post-peak area above the curve reflects stronger glucose recovery relative to total glucose excursion, whereas sustained post-peak elevation produces a lower RBI and corresponds to impaired *β*-cell function. Thus, RBI provides an integrated summary of the balance between recovery and residual glycemic burden within the OGTT profile.

**Figure 3.**
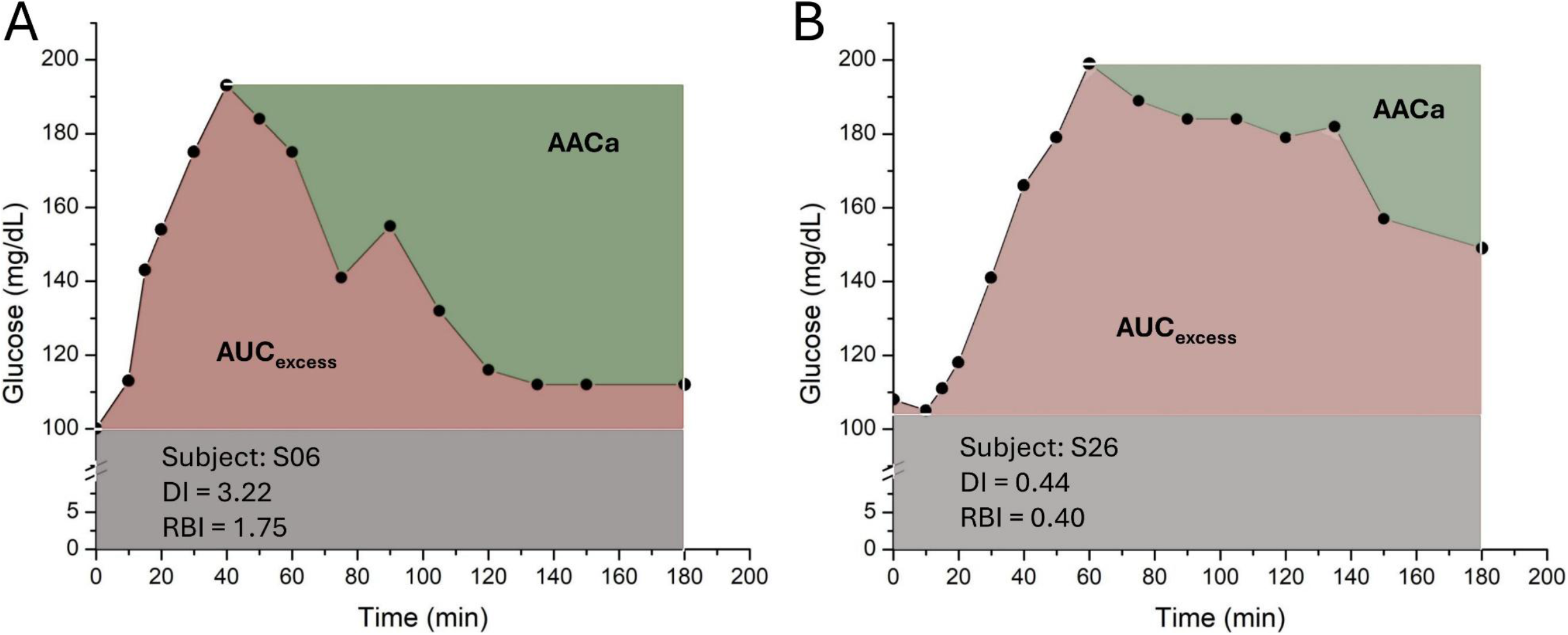
Representative OGTT profiles illustrating recovery-dominant and burden-dominant patterns. Two venous OGTT profiles are shown on the same axis scales. Despite broadly comparable fasting and peak glucose levels, subject S06 showed rapid post-peak recovery, larger AAC_a_, higher RBI, and preserved *β*-cell function, whereas subject S26 showed persistent post-peak hyperglycemia, smaller AAC_a_, lower RBI, and *β*-cell dysfunction. The pink shaded region denotes AUC_excess_, and the green shaded region denotes AAC_a_.

### Reduced sampling preserves RBI performance, but recovery-phase coverage is critical

To assess whether RBI performance required the full densely sampled OGTT profile, we recalculated RBI after downsampling the 0–180 min glucose curve to reduced sampling schedules. The 7-point 0–180 min schedule preserved performance similar to the full profile, with LOO R^2^ of 0.43, Pearson *r* of 0.72, Spearman *ρ* of 0.69, and AUROC of 0.88. More sparse 6-, 5-, and 4-point schedules showed modestly lower continuous DI prediction but retained significant correlations with DI and AUROC values of 0.86–0.89 (Table 2).

**Table 2.** Sensitivity of RBI performance to reduced OGTT sampling density.

| **Metric** | **Timepoints used** | **LOO R<sup>2</sup>** | **OLS R<sup>2</sup>** | **Pearson *r*** | **Pearson *p*** | **Spearman $\rho$** | **Spearman *p*** | **AUROC** |
| --- | --- | --- | --- | --- | --- | --- | --- | --- |
| RBI | 0, 10, 15, 20, 30, 40, 50, 60, 75, 90, 105, 120, 135, 150, 180 | 0.43 | 0.49 | 0.70 | 7.20E-06 | 0.75 | 6.70E-07 | 0.90 |
| RBI 7-point | 0, 15, 30, 60, 90, 120, 180 | 0.43 | 0.52 | 0.72 | 3.20E-06 | 0.69 | 1.50E-05 | 0.88 |
| RBI 6-point | 0, 30, 60, 90, 120, 180 | 0.36 | 0.45 | 0.67 | 2.90E-05 | 0.64 | 7.30E-05 | 0.86 |
| RBI 5-point | 0, 30, 90, 120, 180 | 0.31 | 0.42 | 0.64 | 6.80E-05 | 0.64 | 8.50E-05 | 0.89 |
| RBI 4-point | 0, 30, 120, 180 | 0.34 | 0.43 | 0.66 | 4.20E-05 | 0.70 | 7.10E-06 | 0.89 |

We then tested whether the analyzed OGTT window length influenced RBI performance. RBI calculated over 0–120 min showed lower continuous DI prediction than RBI calculated over 0– 180 min, with LOO R^2^ of 0.12 versus 0.43. Extending the endpoint to 150 min improved performance, and the 0–180 min window showed the strongest performance among baseline-anchored windows. The same pattern was observed for windows beginning at 30 min: RBI30– 120 performed poorly, whereas RBI30–180 performed similarly to the primary RBI analysis, with LOO R^2^ of 0.42 and AUROC of 0.91 (Table 3).

**Table 3.** Sensitivity of RBI performance to OGTT analysis window.

| Start time | End time | LOO R<sup>2</sup> | OLS R<sup>2</sup> | Pearson *r* | Pearson *p* | Spearman $\rho$ | Spearman *p* | AUROC |
| --- | --- | --- | --- | --- | --- | --- | --- | --- |
| 0 min | 120 min | 0.12 | 0.22 | 0.47 | 6.70E-03 | 0.64 | 7.20E-05 | 0.83 |
| 0 min | 150 min | 0.37 | 0.43 | 0.66 | 4.50E-05 | 0.71 | 4.50E-06 | 0.89 |
| 0 min | 180 min | 0.43 | 0.49 | 0.70 | 7.20E-06 | 0.75 | 6.70E-07 | 0.90 |
| 30 min | 120 min | -0.01 | 0.11 | 0.33 | 6.90E-02 | 0.54 | 1.50E-03 | 0.77 |
| 30 min | 150 min | 0.22 | 0.31 | 0.55 | 1.00E-03 | 0.65 | 5.70E-05 | 0.85 |
| 30 min | 180 min | 0.42 | 0.52 | 0.72 | 3.70E-06 | 0.75 | 8.50E-07 | 0.91 |

RBI was recalculated using different OGTT analysis windows to assess the contribution of recovery-phase information beyond the conventional 120-min endpoint. For each window, $G_{\min}$ , $G_{\max}$ , $AUC_{\text{excess}}$ , and $AAC_a$ were recalculated using only glucose values within that interval, and areas were estimated by trapezoidal integration. LOO R<sup>2</sup> was the primary criterion for continuous DI prediction. AUROC was calculated for DI-defined β-cell dysfunction, defined as $DI < 1.58$ , with score direction oriented so that higher AUROC indicates better discrimination. Analysis included 32 participants with measured DI.

These results indicate that dense 15-timepoint sampling is not essential for RBI calculation, but adequate recovery-phase coverage is important. Thus, RBI may be compatible with simplified extended OGTT schedules, whereas conventional 2-hour truncation may limit its ability to estimate DI-associated β-cell compensation.

## Discussion

DI is a physiologically informative measure of *β*-cell function because it evaluates insulin secretion in relation to insulin resistance [4]. However, DI typically requires insulin or C-peptide measurements and model-based estimation, which limits its routine use in clinical and epidemiologic settings. This creates a need for simpler glucose-profile metrics that can capture *β*-cell–relevant information. OGTT provides a dynamic assessment of glucose handling, but its interpretation commonly relies on selected timepoint concentrations or total glucose exposure. These measures are clinically useful, yet they do not explicitly distinguish whether an abnormal glucose excursion reflects a high peak, delayed recovery, or persistent post-peak burden.Because *β*-cell compensation is inherently dynamic, we reasoned that the recovery phase of the OGTT curve may contain information related to DI that is not captured by fasting glucose, 2-hour glucose, HbA1c, or total glucose AUC.

We developed the RBI as a simple geometric measure of post-peak glucose recovery relative to total glucose excursion. RBI showed the strongest prediction of continuous DI among the evaluated single metrics, outperforming fasting plasma glucose, G60, G120, total glucose AUC, and HbA1c. RBI also discriminated DI-defined *β*-cell dysfunction, and representative OGTT profiles illustrated that participants with broadly similar fasting and peak glucose levels can differ markedly in post-peak recovery and RBI. Together, these findings suggest that the balance between glucose recovery and residual glycemic burden is a meaningful feature of the OGTT profile and may provide a compact glucose-only marker related to *β*-cell function.

The physiological rationale for RBI is that post-peak glucose recovery reflects the integrated ability to terminate a glucose excursion. After oral glucose ingestion, the rise and fall of plasma glucose reflect glucose appearance, insulin secretion, insulin sensitivity, and tissue glucose disposal. The recovery phase is especially relevant to *β*-cell compensation because an adequate insulin secretory response, in the context of prevailing insulin resistance, should limit the duration and magnitude of post-peak hyperglycemia. Thus, similar fasting or peak glucose values may have different implications depending on whether glucose declines rapidly or remains elevated after the peak. RBI was designed to quantify this recovery component relative to the overall glucose excursion. A higher RBI reflects stronger post-peak recovery relative to total burden, whereas a lower RBI reflects persistent post-peak elevation and greater residual burden. This interpretation is consistent with the positive association between RBI and DI, but RBI should be viewed as a glucose-profile marker related to *β*-cell compensation rather than a direct measure of *β*-cell function. Unlike DI, RBI does not incorporate insulin or C-peptide secretion, insulin sensitivity, or formal modeling of *β*-cell responsivity; its value is as a compact glucose-only summary of recovery dynamics associated with DI.

Our findings extend prior work showing that dynamic OGTT glucose profiles contain information about underlying metabolic physiology beyond conventional diagnostic thresholds. In the Stanford metabolic phenotyping study, metabolic subphenotypes were predicted using a feature-rich machine-learning framework applied to frequently sampled 16-timepoint OGTT glucose curves [13]. The authors evaluated multiple feature sets within this framework, including two glucose-curve representations. The first used 14 engineered curve features, including timepoint glucose values, AUC-derived measures, peak glucose, curve size, coefficient of variation, time-to-peak, and slope-based features. The second used a two-dimensional reduced representation derived from normalized and smoothed glucose time series. The same framework also evaluated feature sets based on demographics, polygenic risk score, laboratory variables, incretin hormones, HOMA-B, HOMA-IR, and Matsuda index. For *β*-cell dysfunction, the reduced-representation glucose-curve model achieved an AUROC of 0.89, outperforming the non-glucose and surrogate-marker feature sets [13].

RBI differs from these approaches in its simplicity and interpretability. Rather than relying on multivariable feature sets, dimensionality reduction, or machine-learning classification, RBI reduces the OGTT glucose profile to a single geometric recovery-to-burden measure. This construction may explain why RBI outperformed conventional glycemic metrics for continuous DI prediction and also showed strong discrimination of DI-defined *β*-cell dysfunction, with AUROC values of 0.90 for RBI and 0.91 for RBI_30–180_. These discrimination results were comparable to the AUROC of 0.89 reported for the reduced-representation machine-learning model in the source study, while using a single transparent glucose-derived metric. Fasting glucose and HbA1c do not capture the dynamic response to oral glucose; G60 and G120 sample isolated timepoints; and total glucose AUC summarizes exposure without distinguishing whether that exposure reflects early rise, high peak, or delayed post-peak resolution. In contrast, RBI explicitly quantifies post-peak recovery in relation to total glucose excursion, allowing it to capture a DI-relevant feature of the OGTT profile that may be diluted or missed by standard glucose summaries.

RBI may be particularly suitable for CGM-augmented OGTT because its calculation requires sufficient temporal resolution to identify peak glucose and quantify post-peak recovery. Sparse clinical OGTT protocols based on fasting and 2-hour glucose alone cannot capture these geometric features, whereas CGM-augmented OGTT provides dense glucose profiles without repeated venous sampling and may allow practical assessment of recovery-to-burden dynamics. The source study supports this direction by showing that standardized at-home CGM-OGTT curves preserved glucose-response patterns and predicted *β*-cell function using machine-learning models based on glucose-curve features. However, CGM-derived RBI would require separate validation because interstitial glucose differs from venous plasma glucose in sensor accuracy, physiologic lag, and smoothing. Future studies should evaluate agreement between venous and CGM-derived RBI and test whether CGM-derived RBI predicts DI or other *β*-cell function measures. The sensitivity analyses also clarify the practical sampling requirements for RBI. Reduced 0–180 min schedules preserved much of the performance of the full 15-timepoint profile, indicating that dense research-level sampling is not essential. However, truncating the analysis window at 120 min weakened continuous DI prediction, whereas extending the window to 150 or 180 min improved performance. These findings suggest that recovery-phase coverage is more important than maximal sampling density. Therefore, although the 120-min OGTT endpoint is appropriate for diabetes diagnosis, conventional 2-hour OGTT protocols may be less suitable for estimating DI-associated β-cell compensation from glucose-curve geometry.

Several limitations should be considered. First, this was a secondary analysis of a relatively small cohort with measured DI, and the findings require validation in larger and more diverse populations. Second, RBI was evaluated using venous OGTT profiles from a research protocol, and its performance may differ across clinical sampling designs. Although sensitivity analyses suggested that reduced 0–180 min schedules preserved much of the RBI signal, RBI still requires adequate post-peak recovery-phase coverage. If glucose peaks near the final sampled timepoint, subsequent recovery is unobserved and RBI may underestimate recovery. Therefore, RBI is likely better suited to extended OGTT or CGM-augmented OGTT designs than to conventional 2-hour protocols. Third, although DI is a physiologically informative measure of β-cell function, it is itself a derived measure based on C-peptide deconvolution and adjustment for insulin resistance. RBI should therefore be interpreted as a glucose-profile marker associated with DI, not as a direct replacement for insulin- or C-peptide-based assessment of *β*-cell function. Future studies should validate RBI in independent cohorts, define minimal sampling requirements, compare venous and CGM-derived implementations, and evaluate whether RBI adds predictive value beyond established clinical and metabolic markers.

## Conclusion

RBI is a simple glucose-derived index that quantifies post-peak recovery relative to total OGTT glucose excursion. In venous OGTT profiles with measured disposition index, RBI was strongly associated with DI, outperformed conventional glycemic metrics for continuous DI prediction, and discriminated DI-defined β-cell dysfunction. Sensitivity analyses indicated that dense research-level sampling was not essential, because reduced 0–180 min schedules preserved much of the RBI signal. In contrast, truncation at 120 min attenuated performance, supporting the importance of late recovery-phase information. These findings suggest that conventional 2-hour OGTT metrics may be insufficient for estimating DI-associated β-cell compensation from glucose-curve geometry. RBI may therefore provide a transparent and interpretable glucose-profile marker for β-cell–related phenotyping, particularly in simplified extended OGTT or CGM-augmented OGTT settings, although validation in larger independent cohorts is needed.

## Author Contributions

R.Z. conceptualized, designed the study, analyzed the data, wrote the paper, and obtained the funding.

## Data Availability Statement

The OGTT dataset analyzed in this study is publicly available. The Python scripts used for analysis are available from the corresponding author upon reasonable request.

